# Major concerns with the integrity of the mitochondrial ADP/ATP carrier in dodecyl-phosphocholine used for solution NMR studies

**DOI:** 10.1101/329490

**Authors:** Martin S. King, Paul G. Crichton, Jonathan J. Ruprecht, Edmund R.S. Kunji

## Abstract

Hereby, we wish to note our objections to a paper called “Substrate-modulated ADP/ATP-transporter dynamics revealed by NMR relaxation dispersion” by Brüschweiler et al., published in NSMB in 2015^1^. The subject is the yeast mitochondrial ADP/ATP carrier AAC3, which we have studied in great detail ourselves. In particular, we have solved its structure by electron^2^and x-ray crystallography^3^ and have studied its interactions with the specific inhibitors atractyloside (ATR) and carboxyatractyloside (CATR) by single-molecule force spectroscopy^4^. In this paper, the authors claim that AAC3 can be refolded to homogeneity from inclusion bodies produced in *Escherichia coli* by using the detergent dodecyl-phosphocholine (DPC), better known as Foscholine-12 (Anatrace), and that AAC3 is maintained in a folded and active state for the duration of isothermal titration calorimetry (ITC) and NMR experiments. However, in our hands the presence of DPC leads to immediate loss of tertiary structure and inactivation of AAC3 when isolated from the inner membrane of mitochondria, where it was folded and active as shown by functional complementation^2,3^.

To validate the integrity of their refolded protein, the authors first used ITC to measure a Kd of ∼15 µM for the binding of CATR to DPC-refolded AAC3. They then determined CATR-dependent chemical shift perturbations by NMR, giving a Kd of ∼150 µM. No other functional assays were done to verify the activity of the refolded protein, such as transport assays. The inhibitors CATR and ATR differ by one carboxylate group and their affinities have been described in 24 different binding studies (Supplementary Table 1). Yet, the authors only compare their values to a Kd of 192 µM for the binding of ATR taken from one particular study by Babot et al., 2012^6^. Importantly, the µM unit used in this reference was a typographical error and the Kd value has recently been corrected to 192 nM in an erratum^7^. In fact, published Kd values for CATR binding to AAC in the membrane are all in the low nanomolar range (**Supplementary Table 1**), consistent with our own measurements that show that ADP transport by AAC3 is half-maximally inhibited by 1.2 nM CATR (**Fig. 1a**). Thus, the consensus is at least three to four orders of magnitude lower than the Kd values reported for refolded AAC3 in DPC^1^. Our ITC measurements using native AAC3 from yeast mitochondria purified in dodecyl-maltoside/tetraoleoyl cardiolipin gave an average Kd of 72 nM (**Fig. 1b**), which is 200 to 2000-fold lower than those reported in Brüschweiler *et al*.. These experiments were very difficult to do (only 2 out of 10 trials succeeded), as the enthalpic change is low and the apo-state is very unstable in detergent^5^. There are very few polar side chain interactions in AAC3 that stabilize the structure and they are mainly found on the matrix side, where they form intra-domain rather than inter-domain interactions^3^. The ring of transmembrane *α*-helices is held together largely by the lateral pressure of the membrane and counter pressure from the water-filled cavity (**Supplementary Fig. 1a**)^3^, explaining why unliganded AAC3 in detergent micelles is prone to unfolding. Binding of CATR introduces a large number of polar and van der Waals interactions, which cross-link most of the transmembrane α-helices of AAC3 together^3,17^, explaining the high affinity of CATR for AAC3 as well as the improved stability in detergents (**Supplementary Fig. 1b**). The experiments reported here and elsewhere clearly show that the Kd of CATR binding to the folded mitochondrial ADP/ATP carrier is in the low nanomolar range, consistent with its role as a powerful toxin. We further note that to obtain the crystal structure, AAC3 was first inhibited with CATR in the mitochondrial membrane, but then solubilized, purified and crystallized in maltoside detergents in the absence of CATR for 5-7 days^3^. Yet, a clear interpretable density for CATR could be observed^3^, demonstrating that the inhibitor remained bound, consistent with extremely low rates of dissociation. The extremely low affinity of CATR to refolded AAC3 in DPC begs the question whether the binding is specific at all. Only a limited number of chemical shift perturbations for CATR binding to refolded AAC3 are observed^1^ and all of them are dynamic, which is inconsistent with tight binding. Moreover, with few exceptions these residues are not near the known CATR binding site nor are they on structural elements that are involved in CATR binding.

**Figure 1.**
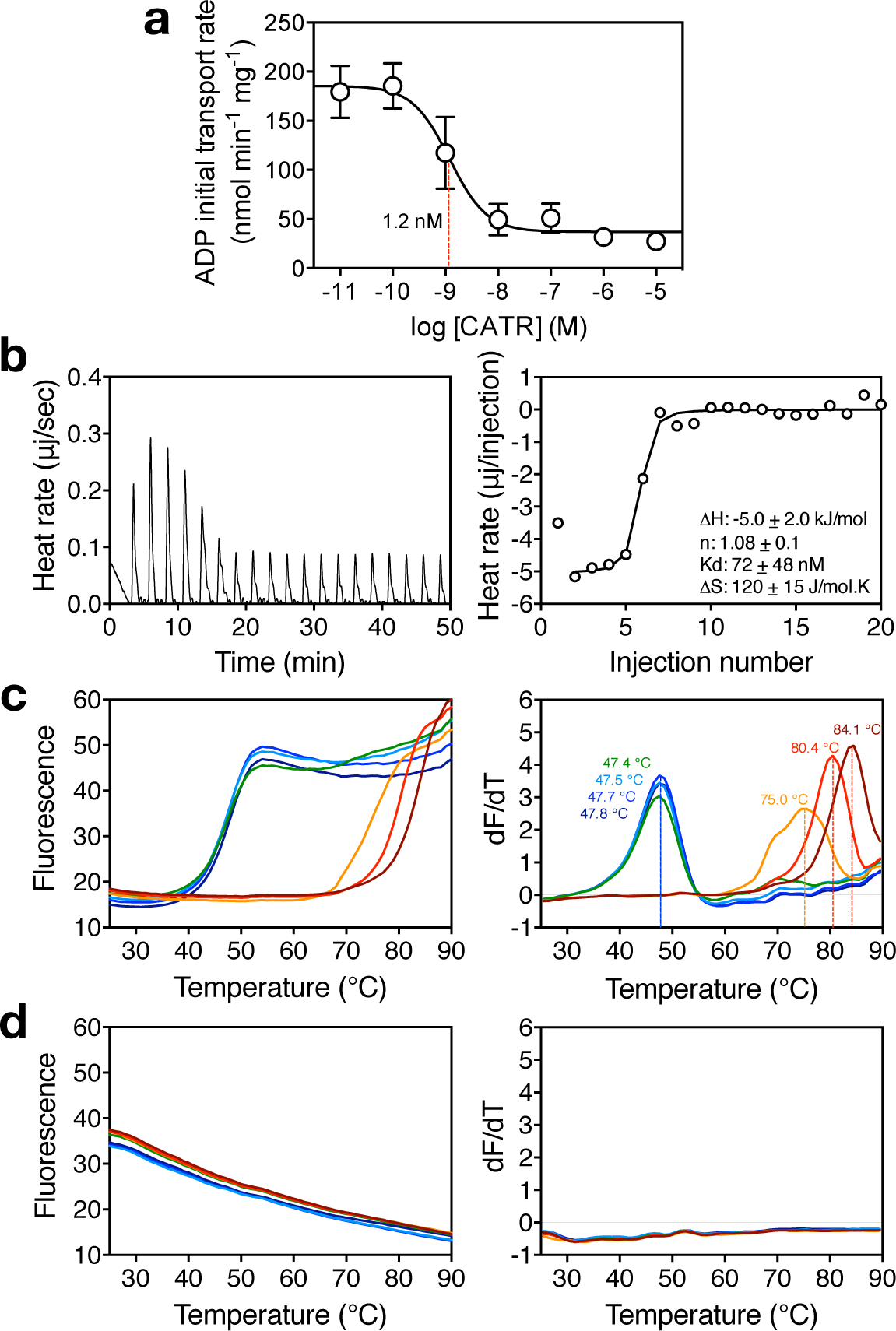
CATR binding and stability of the yeast mitochondrial ADP/ATP carrier AAC3 in dodecyl-maltoside and dodecyl-phosphocholine. (**a**) Inhibition of ADP transport by the yeast mitochondrial ADP/ATP carrier AAC3 by carboxyatractyloside. AAC3 purified from yeast mitochondria in dodecyl-maltoside was reconstituted into liposomes. The CATR concentration required for half-maximal inhibition was determined by measuring the initial ADP uptake rate at different concentrations of CATR (n=3). (**b**) Isothermal titration calorimetry: enthalpy changes associated with the titration of CATR into AAC3 at 10 °C (left panel), and corresponding isotherms fitted to a one site binding model with δH, Kd, stoichiometry and δS as fitting parameters (right panel). (Inset) Average data (± SD) from two titration experiments. Assays were carried out in buffer containing 20 mM PIPES pH 7.0, 100 mM NaCl, 0.1% dodecyl-maltoside, 0.1 mg mL^-1^ tetraoleoyl cardiolipin. (**c**) Thermostability of the yeast mitochondrial ADP/ATP carrier AAC3 diluted into dodecyl-maltoside in the presence of different amounts of CATR (left panel) and derivatives (right panel). (**d**) same as (**c**) but with AAC3 diluted into dodecyl-phosphocholine. The temperature of the protein sample is increased from 25 to 90 °C while protein unfolding is monitored with the fluorophore N-[4-(7-diethylamino-4-methyl-3-coumarinyl)phenyl] maleimide (CPM)^15^. CPM reacts with protein thiols as they become solvent-exposed due to denaturation of the protein to give a fluorescent adduct. The CATR:AAC3 molar ratios were 0 (black), 0.001 (dark blue), 0.01 (light blue), 0.1 (green), 1 (orange), and 10 (red), and 100 (dark red). The protein concentration was approximately 1 μM AAC3.

We also note that the Kd of ADP binding determined by NMR (500 μM)^1^ is ∼85-fold higher than the published consensus values of the carrier in the mitochondrial membrane and ∼25-fold higher than for the solubilized carrier (**Supplementary Table 2**). Residues that were assigned to have chemical shift perturbations induced by ADP are largely on the matrix side of the carrier, far away from the consensus binding site in the central cavity^8-13^. AAC3 has an isoelectric point of 9.82, meaning that it is highly positively charged, whereas CATR and the substrates ADP and ATP are negatively charged molecules at neutral pH. The chemical shift perturbations could represent non-specific interactions of CATR or ADP with AAC3 promoted by the high concentrations and temperatures used in these NMR experiments. We are also concerned about the validity of the dynamic studies measured by NMR relaxation dispersion, as CATR binding should lock the protein in a non-dynamic aborted state, which is why would we could solve its structure by crystallography^3^. We also note that the observed dynamics in the presence of substrate do not provide a plausible structural mechanism for transport.

We have previously demonstrated that DPC is harsh enough to solubilise unfolded mitochondrial carrier protein from *E. coli* inclusion bodies and is able to denature functional well-folded carrier protein prepared in mild non-ionic detergents^14^. In thermostability assays^15^, AAC3 purified from yeast mitochondrial membranes displayed a typical protein melt curve when diluted into the mild detergent dodecyl-maltoside, consistent with thermal denaturation of a folded protein (**Fig. 1c**). When CATR was added at a molar ratio of one or above, a marked shift in stability of AAC3 to higher temperatures is observed, consistent with earlier observations^14,16^. When the same AAC3 preparation was diluted into 3 mM DPC, a high signal was observed with no transition, showing that AAC3 in DPC is in a non-native state (**Fig. 1d**). In this case, CATR addition had no effect, demonstrating that there was no functional binding site. Consistent with these findings, dilution of AAC3 into DPC before reconstitution resulted in a complete loss of CATR-sensitive ADP uptake in liposomes, in contrast to control tests where the protein was diluted into dodecyl-maltoside prior to reconstitution (**Supplementary Fig. 2**). These observations clearly demonstrate that AAC3 is soluble but in a non-native state in DPC. In conclusion, we believe that the data presented by Brüschweiler *et al.* have no biological relevance.

## Acknowledgements

We would like to acknowledge support by the intramural programme MC_UU_00015/1 of the Medical Research Council, UK.

## Supplementary Materials and Methods

### Construction of yeast AAC3 expression strains

The gene coding for AAC3 of *Saccharomyces cerevisiae* was cloned into the yeast expression vector pYES-PMIR2-AAC2 with an N-terminal eight-histidine tag and Factor Xa cleavage site^18^. Expression vectors were transformed by electroporation into *S. cerevisiae* strain WB12 (MATα ade2-1 trp1-1 ura3-1 can1-100 aac1∷LEU2 aac2∷HIS3)^19^, which lacks functional Aac1p and Aac2p carriers. Transformants were selected initially on SC medium minus tryptophan plates, and then on YPG plates, confirming they expressed functional ADP/ATP carriers through complementation.

### Preparation of lipid for protein purification

Tetraoleoyl cardiolipin (18:1) dissolved in chloroform was purchased from Avanti Polar Lipids (Alabaster, Alabama). Typically, 100 mg of lipid was dispensed into a glass vial, and chloroform was removed by evaporation under a stream of nitrogen gas. Lipids were solubilized in 10% (w/v) dodecyl-maltoside by vortexing for 4 h at room temperature to give a 10 mg mL^-1^ lipid in 10% detergent stock. The stocks were snap-frozen and stored in liquid nitrogen.

### Purification of AAC3

A 5-L pre-culture was used to inoculate 50 L of YPG medium in an Applikon 140 Pilot System with an eZ controller. Cells were grown at 30 °C for 72 h, and harvested by centrifugation (4,000 g, 20 min, 4 °C). Mitochondria were prepared with established methods^18^, flash frozen in liquid nitrogen, and stored at −80 °C until use. Yeast mitochondria (1 g total protein) were solubilized in 2% dodecyl-maltoside (Glycon) dissolved in a buffer consisting of 20 mM imidazole, 150 mM NaCl, 20 mM PIPES, pH 7.0, and an EDTA-free complete protease inhibitor tablet (Roche Diagnostics Ltd) by mixing at 4 °C for one hour. Particulate material was removed by ultracentrifugation (140,000 g, 45 min, 4 °C). The soluble fraction was loaded onto a Ni-Sepharose high performance column (Amersham Biosciences) at 1 mL min^-1^ on an ÄKTAprime (GE Healthcare). The column was washed with 40 column volumes of buffer containing 20 mM PIPES pH 7.0, 150 mM NaCl, 20 mM imidazole, 0.1% dodecyl-maltoside, 0.1 mg mL^-1^ tetraoleoyl cardiolipin. The column material was washed with a further 20 column volumes of buffer B containing 20 mM PIPES pH 7.0, 100 mM NaCl, 0.1% dodecyl-maltoside, 0.1 mg mL^-1^ tetraoleoyl cardiolipin. The column material was resuspended with 500 μL buffer B and transferred to a vial containing 5 mM CaCl_2_ and 75 μL Factor Xa (New England BioLabs), vortexed thoroughly, and incubated at 10 °C overnight. The following day, the cleaved protein was separated from the nickel sepharose media using filtration and centrifugation, and the protein concentration determined and the protein snap-frozen and stored in liquid nitrogen.

### Thermostability analysis

Thermostability data were obtained by using the thiol-reactive fluorophore N-[4-(7-diethylamino-4-methyl-3-coumarinyl)phenyl] maleimide (CPM), which undergoes an increase in fluorescence emission following reaction with cysteine residues^20^. A modified procedure using a rotary qPCR machine was used, as described previously^21^. For this purpose, a 5 mg mL^-1^ stock of CPM dissolved in DMSO was diluted 50-fold into buffer containing 20 mM PIPES pH 7.0, 100 mM NaCl, 0.1% dodecyl-maltoside and 0.1 mg mL^-1^ tetraoleoyl cardiolipin, vortexed and the solution was allowed to equilibrate in the dark at room temperature for 10 min. Approximately 1.5 μg of purified protein was added into a final volume of 45 μL buffer containing either 20 mM PIPES pH 7.0, 100 mM NaCl, 0.1% dodecyl-maltoside, 0.1 mg mL^-1^ tetraoleoyl cardiolipin (for the dodecyl-maltoside assays) or 20 mM PIPES pH 7.0, 100 mM NaCl, 3 mM dodecyl-phosphocholine (for the dodecyl-phosphocholine assays), and 5 μL CPM working solution was added, and the solution was vortexed and allowed to equilibrate in the dark for 10 min at room temperature in the presence of increasing concentrations of carboxyatractyloside. Fluorescence of the CPM dye was measured on a Qiagen Rotorgene Q using the HRM channel, which provides excitation light at 440-480 nm with emission detected at 505-515 nm. Measurements were made in 1 °C intervals from 25 – 90 °C with a ‘wait between reading’ set to 4 s, which equated to a ramp rate of 5.6 °C/min, following an initial pre-incubation step of 90 seconds. Data were analyzed and melting temperatures, the peak in the derivative of the melting curve, were determined with the software supplied with the instrument.

### Isothermal titration calorimetry

The enthalpy changes associated with carboxyatractyloside binding to purified AAC3 were recorded with a NanoITC-LV isothermal titration calorimeter (TA Instruments) at 10 °C. Both carboxyatractyloside titrant and protein samples were degassed under vacuum at 10 °C for at least 20 min before the titration experiment. Carboxyatractyloside (500 μM stock prepared in a buffer containing 20 mM PIPES pH 7.0, 100 mM NaCl, 0.1% dodecyl-maltoside and 0.1 mg mL^-1^ tetraoleoyl cardiolipin) was titrated into purified 25 μM AAC3 (0.83 mg mL^-1^ in the same buffer) in 2-μL injections in an initial volume of 170 μL at 3-min intervals with a stirrer speed of 350 rpm. Isotherms were analysed by using the instrument software (NanoAnalyze) and fitted to a one-site binding model with δH, Kd, and stoichiometry as fitting parameters.

### Reconstitution

L-α-phosphatidylcholine and tetraoleoyl cardiolipin were mixed in 20:1 (w/w) ratio (10 mg total lipid) in chloroform and dried under a nitrogen stream, dissolved in methanol, and re-dried to a smear in a 1.5-mL tube. The lipid was emulsified in a buffer solution and solubilised with 55 μL 25% (v/v) pentaethylene glycol monodecyl detergent (Sigma) on ice before addition of 30 μg of purified AAC3 protein to achieve a final buffer composition of 20 mM tris-HCl pH 7.4, 50 mM NaCl and 5 mM ADP (‘internal buffer’), with a volume equivalent to 0.6 mL in the absence of detergent. The purified protein was either added directly to the lipid/detergent reconstitution mix or, for testing the influence of detergents on protein integrity, first diluted 20-fold into 0.1% dodecyl-maltoside or 0.1% dodecyl-phosphocholine and incubated on ice (30 mins) before addition. To form the proteoliposomes, the detergent was stepwise removed from the reconstitution mix through addition of 4 × 30 mg and 4 × 60 mg of Bio-Beads SM-2 in 20-min intervals with gentle mixing in a cold room overnight. The resulting proteoliposome suspension was separated from the biobeads using an empty spin column (Bio-Rad, Hemel Hempstead, UK) and exchanged into ‘external buffer’ (20 mM tris-HCl pH 7.4, 50 mM NaCl, without ADP) using a PD10 desalting column (GE Healthcare, Little Chalfont, UK). The proteoliposomes were eluted in 1.5 mL and diluted a further 9-fold in external buffer before ADP uptake assays.

### ADP uptake assays

Uptake assays were carried out using a Hamilton MicroLab Star robot (Hamilton Robotics Ltd., Birmingham, UK) with 100 µL of the proteoliposome suspension added to each well of a MultiScreenHTS-HA 96-well filter plate (pore size = 0.45 µm Millipore, Billerica, USA), with CATR inhibitor added where required. Uptake of radiolabeled ADP was initiated by the addition of 100 μL of external buffer containing [^14^C]-ADP (2.2 GBq mmol^-1^) to give a final working concentration of 1.5 µM [^14^C]-ADP in the assay. The transport was stopped at 0, 10, 20, 30, 45 s, 1, 2.5, 5, 7.5, 10 and 15 min by the addition of 200 μL ice-cold 20 mM tris-HCl buffer and rapid filtration using a vacuum manifold, followed by an additional wash step with 200 μL ice-cold external buffer and repeated filtration. The amount of [^14^C]-ADP accumulated in the liposomes was measured by the addition of 200 μL MicroScint-20 (Perkin Elmer, Waltham, USA) and quantifying the amount of radioactivity with the TopCount scintillation counter (Perkin Elmer, Waltham, USA). Specific initial uptake rates were calculated using the amount of protein reconstituted into liposomes.

Curves for each uptake were fitted to a ‘plateau and one-phase association’ relationship in Prism (GraphPad). Initial rates were determined from the fit of the entire curve over the 15 min. There is a delay between the addition of radioactive substrate and the washing/vacuuming steps for the zero time point due to the filtration time. For this reason, the fit was not restrained to go through the origin, and the delay was estimated to be 15 s.

## Supplementary figures

**Supplementary Figure 1.**
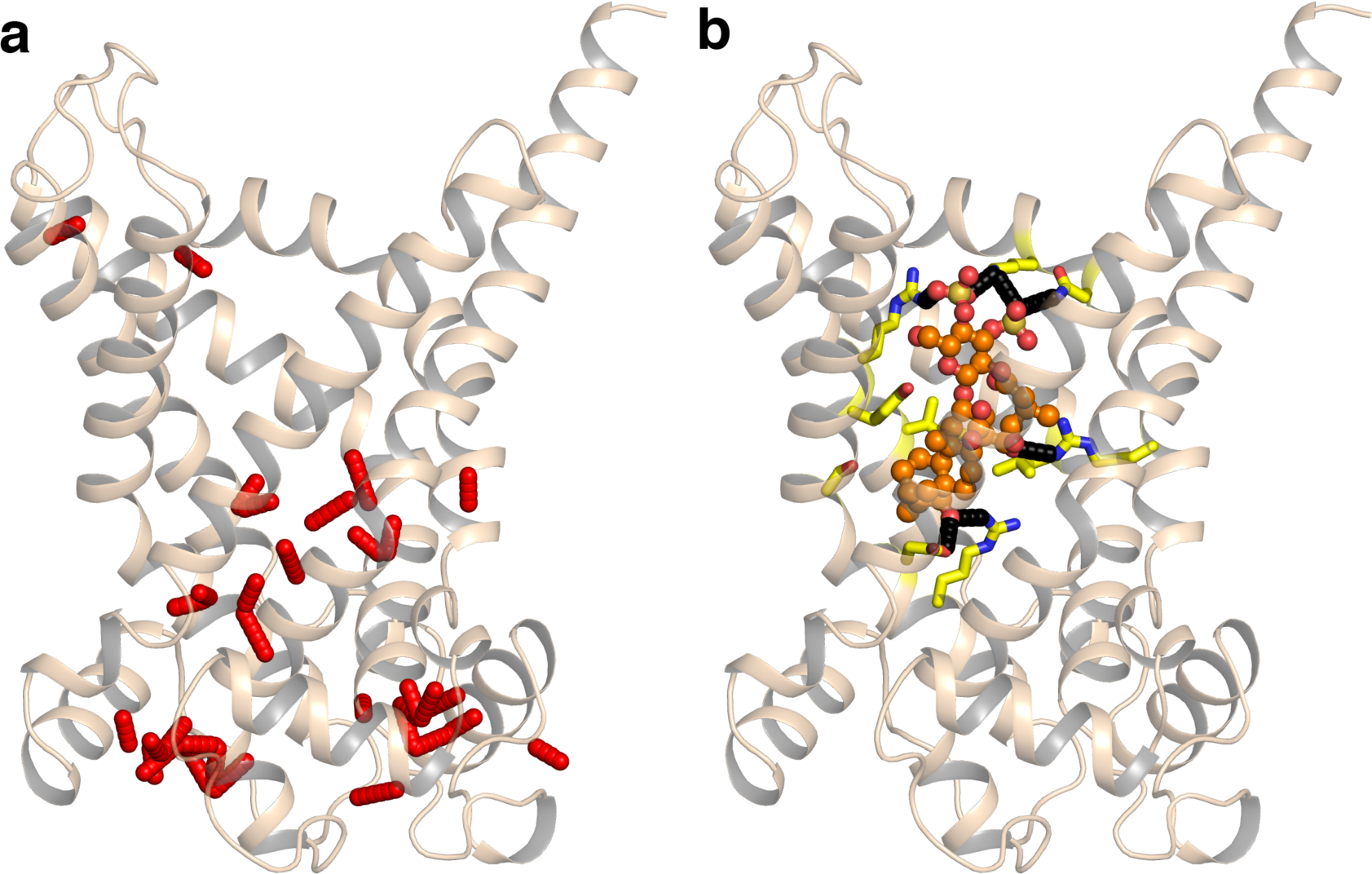
Structural stability of AAC3. (a) The inter-side chain interactions (red sticks) in the structure of AAC3 (cartoon)^22^. There are far fewer polar side-chain interactions between the transmembrane *α*-helices than on the matrix side, and most of them are between residues of the matrix salt bridge network^22,23^ (b) Polar interactions (black sticks) of carboxyatractyloside (orange ball and stick) with AAC3 (cartoon).

**Supplementary Figure 2.**
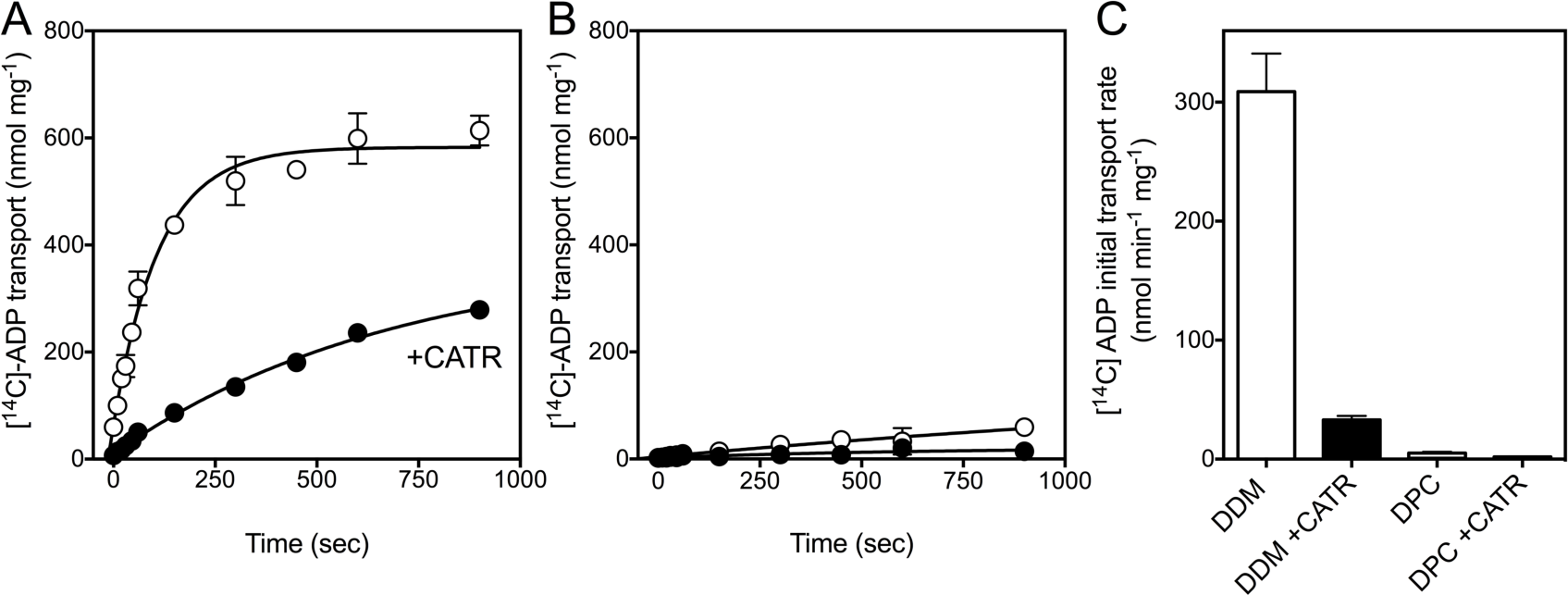
Dodecyl-phosphocholine inactivates the yeast mitochondrial ADP/ATP carrier AAC3. ^14^C-ADP uptake by AAC3 reconstituted into liposomes from (a) dodecyl-maltoside (DDM) or (b) dodecylphosphocholine (DPC). (c) Average initial uptakes rates of experiments in (a) and (b) (n=3). CATR is carboxy-atractyloside.

## Supplementary Tables

**Supplementary Table 1.**
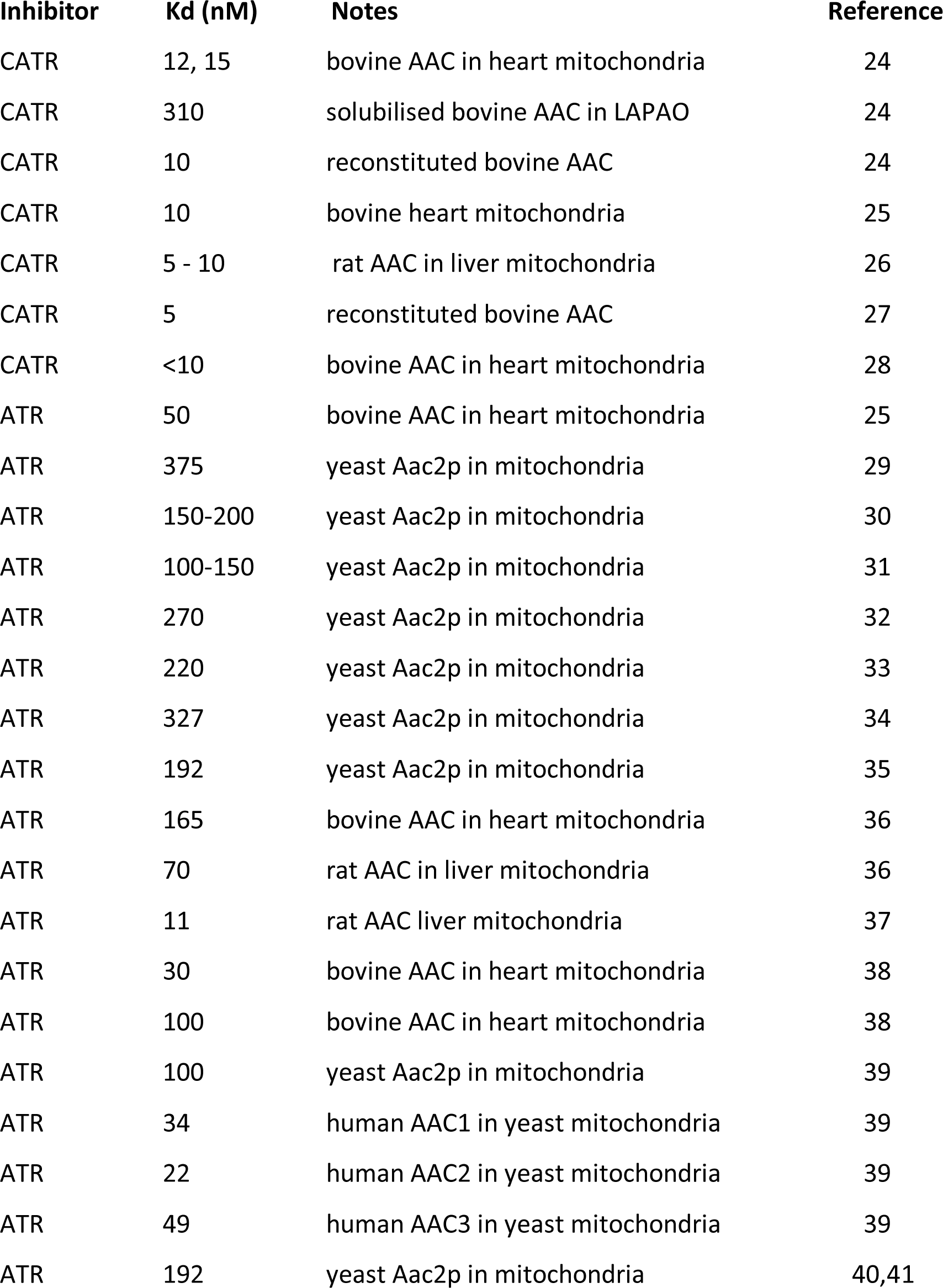
Previously reported values for the dissociation constant (Kd) of carboxyatractyloside (CATR) and atractyloside (ATR) binding to the mitochondrial ADP/ATP carrier

| Inhibitor | K<sub>d</sub> (nM) | Notes | Reference |
| --- | --- | --- | --- |
| CATR | 12, 15 | bovine AAC in heart mitochondria | 24 |
| CATR | 310 | solubilised bovine AAC in LAPAO | 24 |
| CATR | 10 | reconstituted bovine AAC | 24 |
| CATR | 10 | bovine heart mitochondria | 25 |
| CATR | 5 - 10 | rat AAC in liver mitochondria | 26 |
| CATR | 5 | reconstituted bovine AAC | 27 |
| CATR | <10 | bovine AAC in heart mitochondria | 28 |
| ATR | 50 | bovine AAC in heart mitochondria | 25 |
| ATR | 375 | yeast Aac2p in mitochondria | 29 |
| ATR | 150-200 | yeast Aac2p in mitochondria | 30 |
| ATR | 100-150 | yeast Aac2p in mitochondria | 31 |
| ATR | 270 | yeast Aac2p in mitochondria | 32 |
| ATR | 220 | yeast Aac2p in mitochondria | 33 |
| ATR | 327 | yeast Aac2p in mitochondria | 34 |
| ATR | 192 | yeast Aac2p in mitochondria | 35 |
| ATR | 165 | bovine AAC in heart mitochondria | 36 |
| ATR | 70 | rat AAC in liver mitochondria | 36 |
| ATR | 11 | rat AAC liver mitochondria | 37 |
| ATR | 30 | bovine AAC in heart mitochondria | 38 |
| ATR | 100 | bovine AAC in heart mitochondria | 38 |
| ATR | 100 | yeast Aac2p in mitochondria | 39 |
| ATR | 34 | human AAC1 in yeast mitochondria | 39 |
| ATR | 22 | human AAC2 in yeast mitochondria | 39 |
| ATR | 49 | human AAC3 in yeast mitochondria | 39 |
| ATR | 192 | yeast Aac2p in mitochondria | 40,41 |

**Supplementary Table 2:** Previously reported values for the dissociation (Kd) constants of ADP and ATP to the mitochondrial ADP/ATP carrier

| Substrate | K<sub>d</sub> (μM) | Notes | Reference |
| --- | --- | --- | --- |
| ADP | 7 | bovine AAC in heart mitochondria | 24 |
| ADP | 6.6 | bovine AAC in heart mitochondria | 42 |
| ADP | 7 | bovine AAC in heart mitochondria | 43 |
| ADP | 4.1 | rat AAC in heart mitochondria | 43 |
| ADP | 4 | bovine AAC in heart mitochondria | 38 |
| ADP | 2.7 | bovine AAC in heart mitochondria | 44 |
| ADP | 10 | bovine AAC in heart mitochondria | 45 |
| ATP | 12 | bovine AAC in heart mitochondria | 24 |
| ATP | 12 | bovine AAC in heart mitochondria | 43 |
| ATP | 3 | bovine AAC in heart mitochondria | 38 |
| ATP | 5 | bovine AAC in heart mitochondria | 28 |
| ADP | 20 | Solubilised bovine AAC1 | 46 |
| ATP | 20 | Solubilised bovine AAC1 | 46 |

